# Trans effects on gene expression can drive omnigenic inheritance

**DOI:** 10.1101/425108

**Authors:** Xuanyao Liu, Yang I Li, Jonathan K Pritchard

## Abstract

Early genome-wide association studies (GWAS) led to the surprising discovery that, for typical complex traits, the most significant genetic variants contribute only a small fraction of the estimated heritability. Instead, it has become clear that a huge number of common variants, each with tiny effects, explain most of the heritability. Previously, we argued that these patterns conflict with standard conceptual models, and that new models are needed. Here we provide a formal model in which genetic contributions to complex traits can be partitioned into direct effects from core genes, and indirect effects from peripheral genes acting as trans-regulators. We argue that the central importance of peripheral genes is a direct consequence of the large contribution of trans-acting variation to gene expression variation. In particular, we propose that if the core genes for a trait are co-regulated – as seems likely – then the effects of peripheral variation can be amplified by these co-regulated networks such that nearly all of the genetic variance is driven by peripheral genes. Thus our model proposes a framework for understanding key features of the architecture of complex traits.

## 1. Introduction

During the past dozen years, genome-wide association studies (GWAS) have been used to study the genetic basis for a wide variety of complex traits ranging from diseases such as diabetes, Crohn’s disease, and schizophrenia to quantitative traits such as lipid levels, height, and educational attainment [1]. These studies have identified thousands of genetic loci associated with diverse complex traits at genome-wide significance, and in numerous cases it has been possible to dissect the molecular mechanisms that link the identified GWAS variants to disease [2, 3].

Nonetheless, early practitioners of GWAS were surprised to find that even the strongest GWAS hits tend to have modest effect sizes on risk, and that all the genome-wide significant hits in combination explained only a small fraction of the expected genetic component of risk [4]. For example, the 18 genome-wide significant loci for type 2 diabetes identified by 2010 explained just 6% of the expected heritability; for height, the 40 genome-wide significant loci explained just 5% of the heritability [4]. Over time, estimates of explained heritability have only increased modestly, even with much larger sample sizes and many more significant loci [5]. This initial observation that genome-wide significant loci only capture a small proportion of the expected genetic heritability became known as the problem of “missing heritability”. Subsequent work has largely resolved this initial mystery by showing that most of the missing heritability is due to large numbers of small-effect common variants that are not significant at current sample sizes [6, 7, 8, 5].

While this initial mystery has been resolved, the resolution led to another surprising finding – the large numbers of small-effect variants tend to be spread extremely widely across the genome and implicate a considerable fraction of all genes expressed in relevant tissues. Indeed, for many traits, a large fraction of the genome contributes to heritability [6]. For example, between 71–100% of 1MB windows in the genome are estimated to contribute to the heritability of schizophrenia [8]. Similarly, a recent study of polygenic prediction models found that for most of the diseases studied, the models achieved peak accuracies when assuming that 0.1%-1% of SNPs have causal effects [9].

We recently argued that the data suggest that a large fraction of all genes expressed in relevant tissues can affect a phenotype, and that much of the trait variance is mediated through genes that are not tightly involved in the trait in question [10]. These observations appear at odds with conventional ways of understanding the links from genotype to phenotype. Much of the progress in classical genetics has come from detailed molecular work to dissect the biological mechanisms of individual mutations. That type of work is predicated on the expectation that there should be a relatively direct molecular pathway from genotype to phenotype. Yet the situation for complex traits seems quite different, and thus it remains unclear how we should understand the molecular mapping from genotype to phenotype.

Specifically, the data suggest several key questions:

- *Why does such a large portion of the genome contribute to heritability?*
- *Why do the lead hits for a typical trait contribute so little to heritability?*
- *What factors determine the effect sizes of SNPs on traits?*

In this paper, we develop a statistical model to explore these questions. Our model necessarily simplifies a more complex reality, and elides specific details of biology and genetic architecture that vary across traits. Nonetheless, we believe it is essential for the field to develop conceptual models for understanding complex trait architecture, and the model proposed here is a step in that direction.

The central thesis of the present paper is that known properties of cis- and trans-regulatory effects(i.e., cis and trans expression- or protein-QTLs) provide essential clues to understanding key features of the architecture of complex traits.

### Key observations

As reviewed in our previous paper [10], a conceptual model of complex traits should allow for the following observations:

1. The most important loci contribute only a modest fraction of the total heritability [5]. Nevertheless, for many traits the most significant signals are located near genes that make functional sense. This has been established both by detailed molecular dissection of top hits as well as by enrichment analyses of significant loci (although the strength of enrichment is generally modest and varies among traits) [11, 12, 13, 14].
2. The bulk of the heritability can be attributed to a huge number of common variants with very small effect sizes. Moreover, these variants tend to be spread very broadly across the genome [8]. For traits such as schizophrenia and height, analyses suggest that up to half of all SNPs may be in linkage disequilibrium with causal variants [10].
3. Consistent with the latter observations, genes with putatively relevant functions (e.g., neuronal functions for schizophrenia and immune functions for Crohn’s disease) contribute only slightly more to the heritability than do random genes, as measured on a per-SNP basis. (While gene functional annotations are imperfect, it is worth noting that other kinds of experiments, such as genome-scale CRISPR-screens, often yield much stronger functional enrichments than seen in most GWAS data [15, 16, 17].) The clearest functional pattern is that genes not expressed in relevant cell types do not contribute significantly to heritability [10].
4. Similarly, the per-SNP heritability in tissue-specific regulatory elements is only modestly increased relative to SNPs in broadly active regulatory elements, provided that they are active in relevant tissues [10]. Thus, various lines of evidence indicate that the heritability of a typical complex trait is driven by variation in a large number of regulatory elements and genes, spread widely across the genome and mediated through a wide range of gene functional categories.
5. For most complex traits, the heritability is dominated by common variants [5, 7, 18]. While rare variants with large effect sizes do exist for some complex traits (and often highlight genes with key biological roles, e.g., [19, 20]), rare variants are generally not major contributors to the overall phenotypic variance.
6. Protein-coding variants typically contribute very little to complex disease risk (*~*10% [21, 22]). The SNPs that contribute to heritability are highly enriched in noncoding regions, including especially in active chromatin regions [21, 22, 23]. There is strong enrichment of cis and trans eQTLs among GWAS hits (albeit still a considerable gap in linking all hits to eQTLs) [24, 25, 26, 27].

Together, these points suggest an architecture in which some genes (and their regulatory networks) are functionally proximate to disease risk. These genes tend to produce the biggest signals in common-and rare-variant association studies, and they tend to be the most illuminating from the point of view of understanding disease etiology. However, they are responsible for only a small fraction of the population variance in disease risk. This implies that the bulk of the heritability is explained by genes that have a wide variety of functions, many of which have no obvious functional connection to disease aside from being expressed in disease-relevant tissues. Lastly, most of the GWAS hits are in noncoding, putatively regulatory regions of the genome, indicating that the primary links between genetic variation and complex disease are via gene regulation.

## 2. The omnigenic model

We previously proposed the omnigenic model as a conceptual framework to explain the observations above (Figure 1) [10, 28]. The omnigenic model partitions genes into core genes and peripheral genes. Core genes can affect disease risk directly, while peripheral genes can only affect risk indirectly through transregulatory effects on core genes. Two key proposals of the omnigenic model were that (1) most, if not all, genes expressed in trait-relevant cells have the potential to affect core gene regulation, and (2) that for typical traits, nearly all of the heritability is determined by variation near peripheral genes. Thus, while core genes are the key drivers of disease, it is the cumulative effect of many peripheral gene variants that dictates polygenic risk^1,2^.

**Figure 1:**
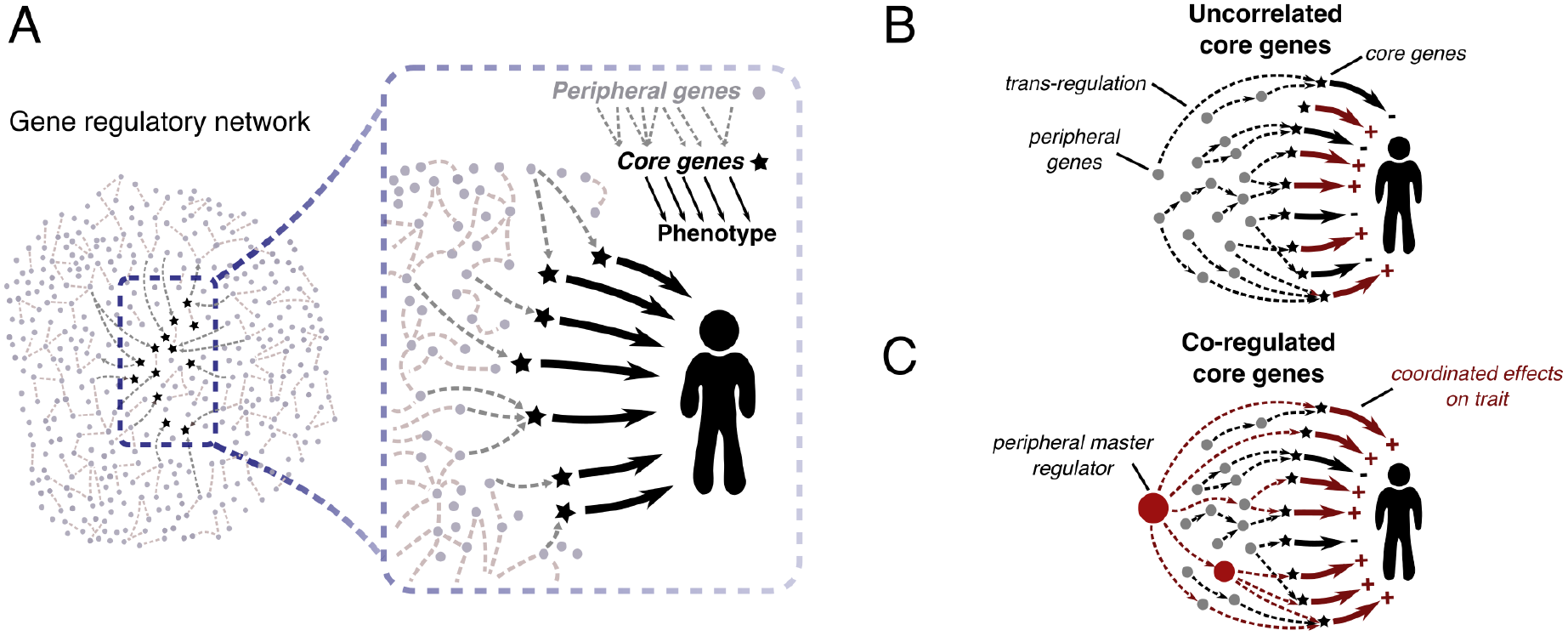
Our model starts by defining “core” genes as the set of genes that exert direct effects on a trait: i.e., not mediated through regulation of other genes. **(A)** Core genes are embedded in gene regulatory networks, such that regulatory effects from all other expressed genes (i.e., peripheral genes) may affect core gene regulation and thus affect the trait indirectly. Most heritability is due to variation at peripheral genes. **(B)** According to the model, most cis-regulatory variants for peripheral genes are also weak trans-QTLs for core genes, and the direction of effect varies across core genes. Thus, typical peripheral variants make tiny contributions to heritability, but because there are so many, they are responsible for most of the heritability. **(C)** Some peripheral genes drive coordinated regulation of multiple core genes with shared directional effects, and can thus stand out as relatively strong GWAS hits. As discussed later in the paper, likely examples include KLF14 and IRX3/5 [2, 31].

### Definitions

We define a gene as a **core gene** if and only if the gene product (protein, or RNA for a noncoding gene) has a direct effect–not mediated through gene regulatory networks–on cellular and organismal processes leading to a change in the expected value of a particular phenotype. All other genes expressed in relevant cell types are considered **peripheral genes**, and can only affect the phenotype indirectly through regulatory effects on core genes. Third, **unexpressed genes**, i.e., genes that are not expressed in trait-relevant tissues, are assumed not to contribute to heritability.

Importantly, this definition of core genes implies that the phenotype of an individual is conditionally independent of the peripheral genes, given the expression levels and coding sequences of the core genes^3,4^ (Figure 2).

**Figure 2:**
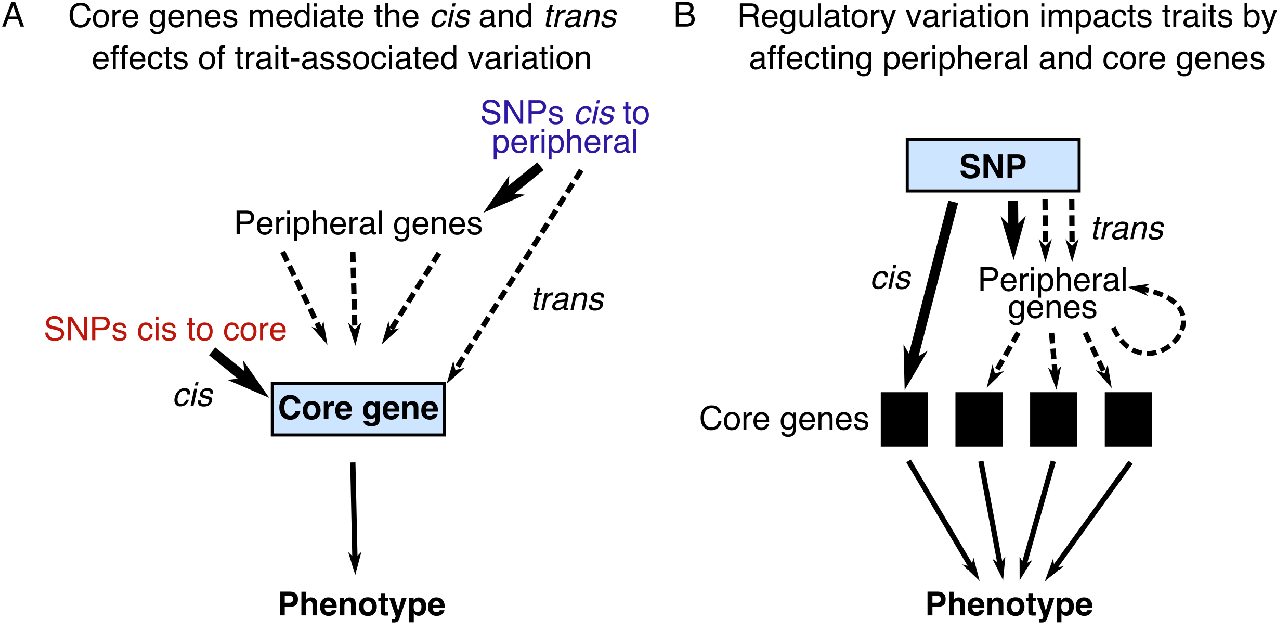
Causal pathways for variants affecting a trait through core genes. By definition only the core genes exert direct effects on the phenotype. We assume that they do so mainly through variation in expression levels. **(A)** Cis and trans regulatory effects are funneled through core genes to affect the phenotype. **(B)** From the vantage point of a regulatory QTL SNP, effects fan out through cellular regulatory networks to affect one or more core genes.

Most peripheral genes make relatively small contributions to heritability. However, some peripheral genes, such as transcription factors and protein regulators, play important roles because they are key regulators of multiple core genes (Figure 1C). As discussed below, when a single peripheral gene coordinately regulates multiple downstream core genes with shared directions of effect, there is a potential for relatively large effect sizes at that peripheral gene. We refer to such genes as **peripheral master regulators**. Selective constraint on master regulators may be particularly strong, with the result that GWAS signals at these loci are often smaller than might be expected from their intrinsic importance [32].

In the Discussion section of this paper, we provide examples to illustrate these definitions.

### A quantitative phenotype model based on core gene expression

To model the contribution of core and peripheral genes to complex trait heritability, we now propose a quantitative model that links phenotypic variation to the expression levels of core genes in a disease-relevant tissue:

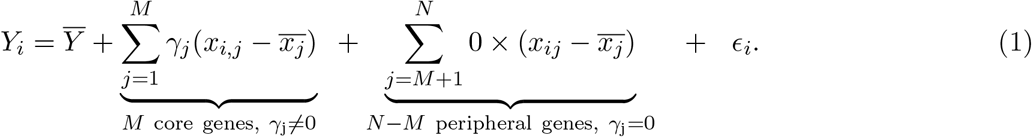

Here *Y_i_* denotes the phenotype value in individual *i* and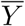 is the population mean phenotype. *γ_j_* denotes the *direct* effect of a unit change in expression of core gene *j* on E(*Y_i_*); and *x_ij_* is the expression of gene *j* in individual *i* (with population mean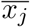). There are *M* core genes, out of *N* total expressed genes. The error term *∈_i_* represents random effects and is assumed to be independent of genotype and gene expression^5,6^.

Importantly, this model assumes that each core gene affects the expected phenotype value as a linear function of its expression level (with slope *γ_j_*). Notice that the expression levels of peripheral genes do not have *direct* effects on the phenotype *Y*, but may affect *Y indirectly* by modifying the expression of core genes as trans-QTLs. This model assumes the simplest possible relationship between expression levels of core genes and the phenotype: namely that the expression of each core gene is linearly related to the expected phenotype value and with no additional interaction terms. One might reasonably argue that biological reality is more complex than assumed here, however simple models such as these can be particularly useful for elucidating general principles.

### Core and peripheral genes; cis and trans eQTLs

These definitions imply a close connection between the core-peripheral distinction and cis vs. trans eQTLs (or pQTLs). Genetic variants that are cis-eQTLs to core genes can affect disease risk directly through their effects on core gene expression. In contrast, genetic variants elsewhere in the genome can only affect core gene expression as trans-eQTLs, presumably mediated through peripheral genes (Figure 2). As discussed in the next sections, trans-eQTLs generally have very small effect sizes relative to cis-eQTLs. Thus, each peripheral variant is likely to have small effects on disease risk, except for those that regulate multiple core genes with consistent directions of effect.

In the next sections we explore the implications of this model from two viewpoints. First, we focus on the combination of cis and trans effects converging onto a core gene. Second we explore the properties of SNP effects, which may fan out to impact expression of multiple core genes, and thereby disease risk.

#### Optional Box 1.

Incorporating genetic variation into the model. Suppose that there are *S_j_* distinct eQTLs for core gene *j*, of which *S_j_*,_cis_ are in cis, and *S_j_ − S_j_*,_cis_ are in trans. Let *β_s,j_* denote the effect size of eQTL *s* ∈ 1 & *S_j_* on expression of gene *j* (each additional copy of the alternate allele at *s* increases expression of *j* by *β_s,j_* units). Then, assuming a linear model of eQTL effects, the expected expression level of gene *j* in individual *i*, relative to the population mean, depends on that individual’s genotype as follows:

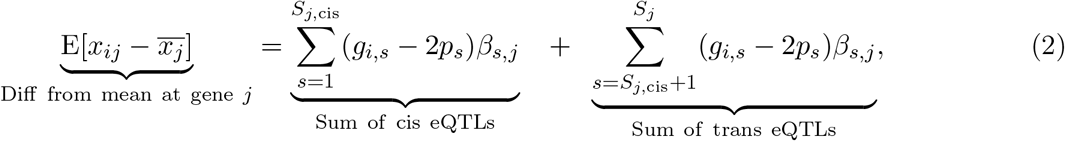

where *g_i,s_* ∈ {0, 1, 2*}* is the genotype of individual *i* at SNP *s*, and *p_s_* is the population allele frequency at SNP *s* (this factor reflects the average genotype). Then plugging Eq. 2 into Eq. 1 and assuming no interaction effects, we can write the expected phenotype *Y* for individual *i* in terms of their genotype:

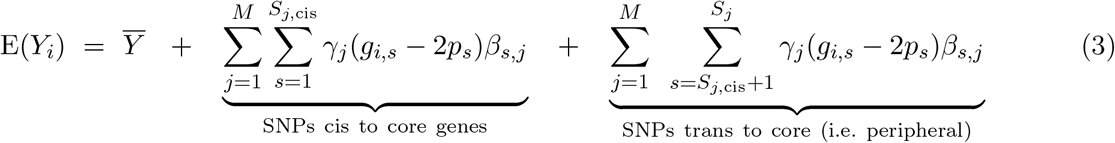

In this last expression, SNPs or other variants near to core genes affect their expression as cis-eQTLs, and SNPs elsewhere in the genome act as trans-eQTLs. These genetic effects on core gene expression, in turn, change the expected phenotype value *Y_i_* by the factor *γ_j_* for each core gene *j*. The form of Equation 3 is reminiscent of a polygenic risk score, except that in a polygenic score, the terms are collapsed into a single value per SNP because we do not currently know the identities of the core genes, nor the γs or βs.

## 3. Core gene effects on heritability

A key hypothesis of the omnigenic model is that most of the heritability for complex traits comes from peripheral genes. We can now use this model, combined with existing data about the genetic architecture of gene expression, to better understand why this may be.

### The heritability of expression is dominated by many small trans effects

The first key question is: for a typical gene, what fraction of heritability of gene expression is determined in cis vs. trans? This is a difficult quantity to measure as most studies are underpowered to detect trans eQTLs, and thus estimates of trans heritability must rely on statistical methods that aggregate weak signals. However, the literature is reassuringly consistent across a range of study-types, indicating that around 60-90% of genetic variance in expression is due to trans-acting variation (Table 1)^7^. For clarity we will refer to the fraction of trans heritability as 70%, while noting uncertainty in the precise value.

**Table 1:** Studies of cis vs trans heritability. Despite some variability across species, cell types, and analytic methods, these studies all indicate that most heritability of gene expression is due to trans variation. Data refer to mRNA expression, except the last two rows which are for protein expression. See the Supplement for further notes on these studies.

| Percent $h^2$ in trans | Tissue/ organism | Platform | Sample size | Method | Reference |
| --- | --- | --- | --- | --- | --- |
| **88%** | LCL from admixed inds | Affymetrix Array | 89 | African-European ancestry | Price 2008 [33] |
| **76%, 61%** | Drosophila, whole body | RNA-seq | multi-fly pools | fly hybrids | McManus 2010 [34] |
| **76%, 63%** | adipose, blood | custom array | 638, 687 | cis/trans IBD in families | Price 2011 [35] |
| **70%, 65%, 64%** | adipose, LCL, skin | Illumina Array | 856 | twin design | Grundberg 2012 [36] |
| **77%, 69%** | peripheral blood | Affymetrix Array | 2,752 | twin design, LD Score | Wright 2014 [37, 38] |
| **72%** | yeast segregants | RNA-seq | 1012 | cis vs. trans eQTLs | Albert 2018 [39] |
| **62%** | mouse liver | RNA-seq | 192 | GCTA | This study; data [40] |
| **72%** | mouse liver (proteins) | Mass Spec | 192 | GCTA | This study; data [40] |
| **78%** | human plasma (proteins) | protein aptamers | 3301 | LD Score Regression | This study; data [41] |

Despite the overall importance of trans effects, trans-eQTLs are notoriously difficult to find in humans [43, 25, 42, 44]. This is partly due to the extra multiple testing burden on trans-eQTLs, but is mainly due to the small effect sizes of trans-eQTLs. To illustrate this, Figure 3 plots the cumulative distributions of cis- and trans-effects in a sample of 913 individuals in whole blood, showing that trans effects are uniformly small compared to cis effects, with only a handful reaching significance. Given that most trans-eQTLs are far below the detection threshold for current eQTL studies it is difficult at present to estimate how *many* trans-eQTLs act on a typical gene. Nonetheless, since *~*70% of the heritability of expression is in trans, this implies that typical genes must have very large numbers of weak trans-eQTLs.

**Figure 3:**
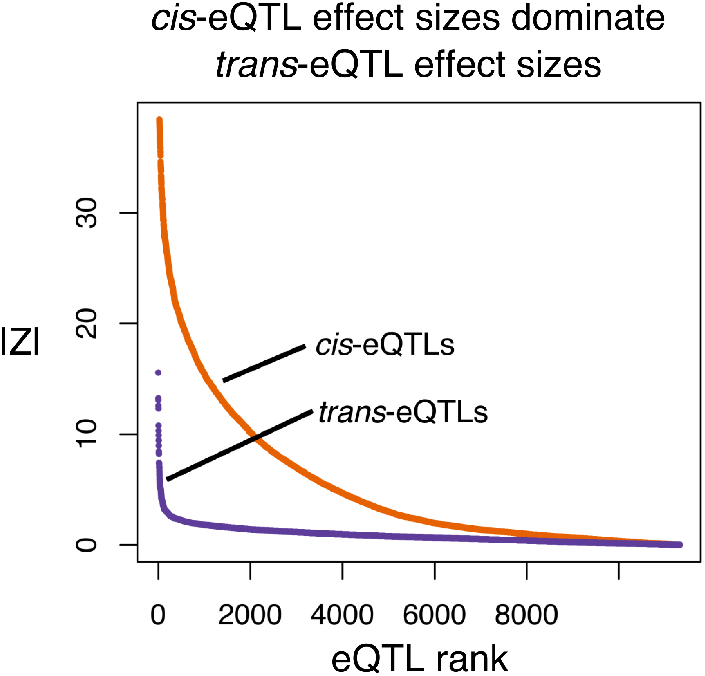
Cumulative distributions of signal sizes for the strongest cis and trans eQTLs.

for each expressed gene in whole blood (n=913). The signals are plotted as |Z|-scores; note that Z^2^ is proportional to the genetic variance contributed by each SNP. To reduce the biasing effects of winner’s curse and the very different numbers of tests in cis and trans, we first identified the most significant cis- and most significant trans-signal for every gene in one data set [37] and plot here the distribution of |Z|-scores for those SNP-gene pairs in a replication data set [42].

If we assume that typical complex traits have, perhaps, hundreds of core genes, and that each is likely affected by many weak trans eQTLs, this starts to explain why so much of the genome contributes heritability for typical traits.

### Core and peripheral contributions to heritability

We next use this model to explore how much of the heritability is determined by cis-regulatory effects on core genes vs. trans-regulatory effects from peripheral genes.

Eq. 1 models the relationship between the phenotype value *Y* and the expression of the core genes. From this, we can compute Var(*Y_i_*) in terms of genetic variances and covariances of gene expression of core genes (see Box 2 for details):

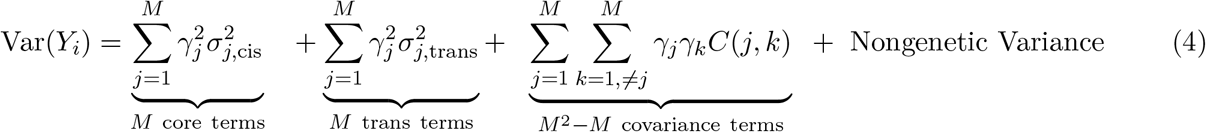

where *σ*^2^_*j*,cis_ and *σ*^2^_*j,trans*_ are the cis and trans genetic variances underlying expression of gene *j*, and *C*(*j, k*) denotes the genetic covariance of expression of genes *j* and *k*. Apart from the special case of core genes that are adjacent in the genome, genetic covariances of expression must be determined by trans effects. As before, *γ_j_* measures the effect of a unit change in expression of gene *j* on the phenotype *Y*.

To interpret Eq. 4 we consider two main cases^8^ depending on the sum of covariance terms *γ_j_ γ_k_C*(*j, k*). These two scenarios predict that between *~*70% to nearly all of the heritability is likely determined by trans effects (Figure 4).

**Figure 4:**
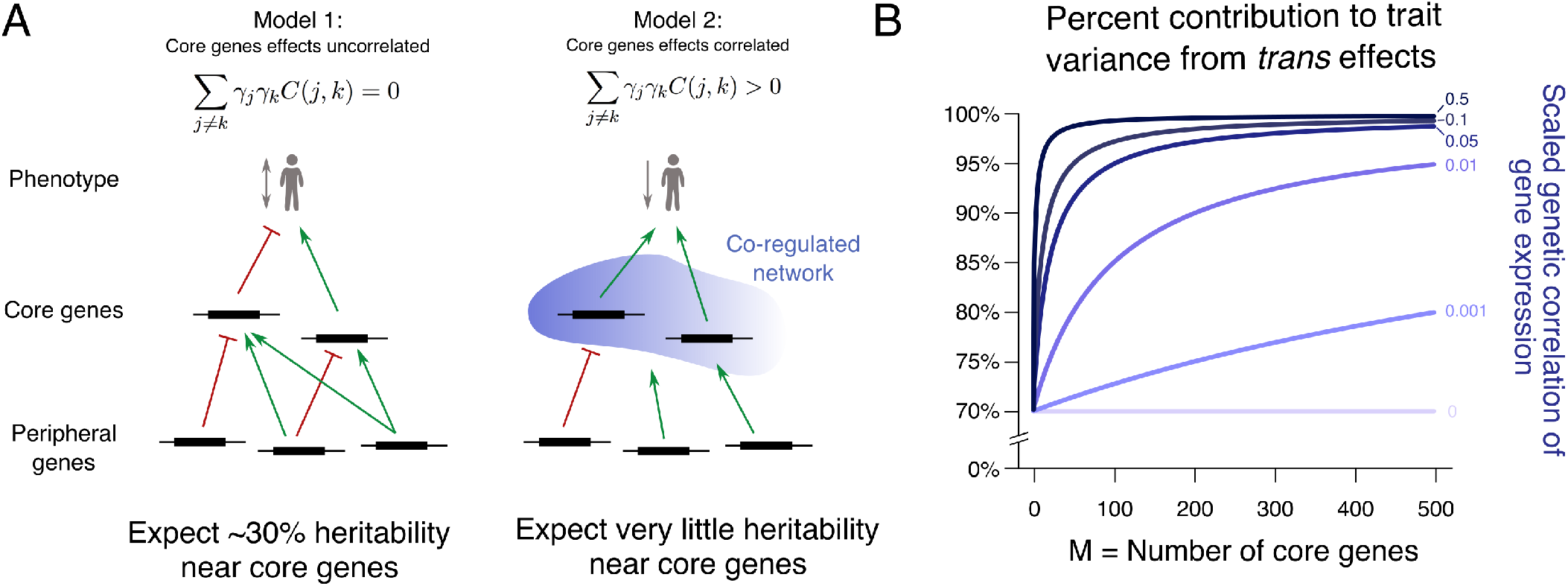
Modeling predicts that from 70% to nearly 100% of heritability is driven by weak trans effects. **(A)** In Model 1 we assume that expression of core genes tends to be relatively independent. In this case we predict that about 30% of heritability is in cis to the core genes. In Model 2 we assume that core genes are often co-regulated, with coordinated directions of effects. In this case, for any given individual, the aggregated effects of peripheral variants are partially shared across core genes, while the directions of ciseffects at core genes may be up, or down, independently across genes. This effectively transfers most of the heritability out to a large number of peripheral regulators. **(B)** Illustration of the fraction of genetic variance due to trans variance and covariance effects (Eq. 4). (Simplifications for plotting: σ and | γ | constant; the “scaled correlation” is *E[sign*(γ_i_ γ_j_) × ρ(j,k)] where ρ(j,k) = C(j,k)/σ_j_σ_k_ is the genetic correlation of genes j and k).

### Model 1: Core genes generally not co-regulated

Suppose that core genes tend to be dispersed in gene regulatory networks, or that the signs of their effects on disease are not coordinated. Specifically, we assume that the average value of *γ_j_ γ_k_C*(*j, k*), computed across pairs of core genes, is approximately 0. In this case, we can ignore the sum of covariance terms.

Now, the fraction of the genetic variance that comes from regulatory variants cis to core genes is simply the average fraction of cis heritability in core genes. Assuming that core genes are typical of genes overall, we can predict that about 30% of heritability comes from cis-regulatory variants acting on core genes, and 70% from trans effects, mainly from peripheral genes^9,10^.

### Model 2: Core genes generally co-regulated

Crucially, there are nearly *M*-fold as many covariance terms in Equation 4 as variance terms. Hence, if a considerable fraction of core genes are either co-regulated with shared directions of effects, or negatively co-regulated with opposite directions of effects–(i.e., *γ*_*j*_ *γ_k_C*(*j,k*) *>* 0)–then the sum of covariance terms can dominate the genetic variance for trait *Y*. Since covariances are primarily driven by trans effects, co-regulated networks could potentially act as strong amplifiers for trans-acting variants that are shared among core genes in those networks.

For example, a recent paper by Gandal et al. identified several co-expressed gene modules that are either up-regulated or down-regulated in various psychiatric conditions, compared to controls [45]. We hypothesize that such modules may often contain multiple core genes with covarying directions of effects, as well as genetic co-regulation. If this is the case, then most of the phenotypic variance may be driven by (trans-acting) covariance terms.

There has been little work so far on measuring the genetic basis of gene expression correlations. Nonetheless, the work to date shows that expression covariance is substantially driven by genetic factors. For example, Goldinger et al. (2013) studied heritability of principal components in a data set of whole blood gene expression from 335 individuals [46]. They concluded that there was a strong genetic component in the lead PCs, with an average heritability of 0.39 for the first 50 PCs.

Similarly, Lukowski et al. (2017) tested for genetic covariance between gene pairs, and identified 15,000 gene pairs (0.5% of all gene pairs) with significantly nonzero genetic covariance at 5% FDR [47]. Since the significance test is likely underpowered, there are probably many more gene pairs with covariance. For example, for the 10% of gene pairs with the highest phenotypic correlation, the average genetic correlation is 0.12 (Supplementary Information). This magnitude is potentially large enough to make an important contribution to heritability (Figure 4B). However, their data show roughly equal numbers of positive and negative genetic correlations overall. Since the overall contribution of the covariance terms depends on the average of *γ_j_ γ_k_C*(*j,k*), this means that in order for the covariance terms to matter, either core gene pairs would have to be enriched for positive covariances, or the sign of the covariance for a given pair would have to tend to match the sign of *γ_i_ γ_j_*. Both of these scenarios seem plausible but will require further study.

In summary, each core gene is likely affected by large numbers of weak trans-acting (peripheral) variants. Assuming that a typical trait might have hundreds of core genes, this may help to explain why so much of the genome contributes to heritability for typical traits. Furthermore, this model suggests that most trait heritability is mediated through trans effects, especially if core genes tend to be positively co-regulated.

#### Optional Box 2.

Cis and trans contributions to heritability. Eq. 1 models the relationship between the phenotype value *Y* and the expression of the core genes. Then, using standard rules of probability, the phenotypic variance is given by

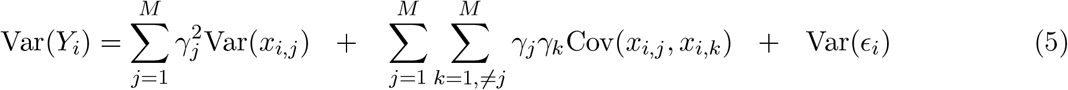

where the variances and covariances are computed across individuals (subscript *i*). Here, the first sum adds up the variances of expression of the core genes, and the second sum adds the covariances of expression between all pairs of core genes. Var(*∈_i_*) encompasses nongenetic effects and is not relevant for understanding heritability.

To interpret these, we need to write the expression variances and covariances in terms of genetic contributions. As before, we assume fully additive models, no interaction terms, and in this case linkage equilibrium between eQTLs:

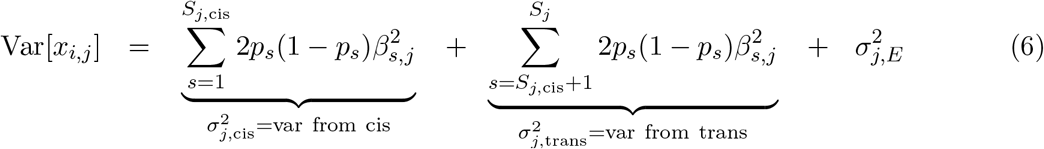

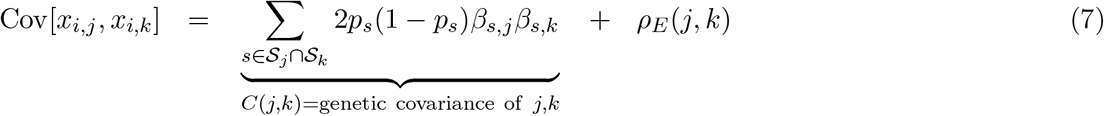

The equations above define the cis and trans components of expression variance of gene *j* (*σ*^2^_*j*,cis_ and *σ*^2^_*j,trans*_, respectively), and the genetic covariance of genes *j* and *k* (*C*(*j,k*)). Here *σ*^2^_*j,E*_ and *ρ_E_*(*j,k*) are the nongenetic variance and covariance respectively. *S_j_* and *S_k_* denote the sets of eQTLs for genes *j* and *k* respectively. Now we can decompose the additive genetic contributions to Var(*Y_i_*) as follows:

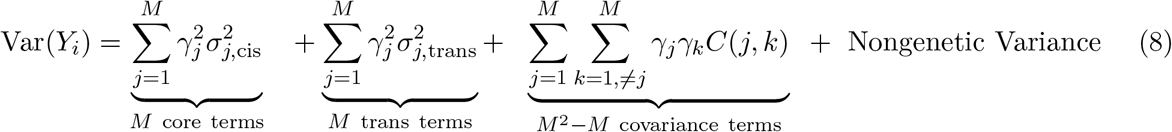

where the nongenetic variance consists of Var(*∈_i_*) plus sums of *σ*^2^_*j,E*_ and *ρ_E_*(*j,k*).

Equation 8 suggests why so much of the genetic basis of complex traits come from trans (mainly peripheral) effects. Since there are so many covariance terms, and since these are mainly driven by trans effects, these can dominate the genetic variance in *Y* if the average value of *γ _j_ γ_k_C*(*j,k*) is non-negligible compared to the average of γ^2^_*j*__σ_^2^_*j*_.

## 4. SNP effect sizes on disease risk

In the previous section we focused on the behavior of the model from the point of view of core genes–which collect QTL effects from cis and trans variants. We now turn our attention to a SNP-centric viewpoint. The effects of a single SNP potentially fan through multiple core genes to affect the phenotype (Figure 2B, Figure 5). The SNP effect sizes that are measured in GWAS correspond to the aggregated effects of each SNP on all core genes, as described next.

**Figure 5:**
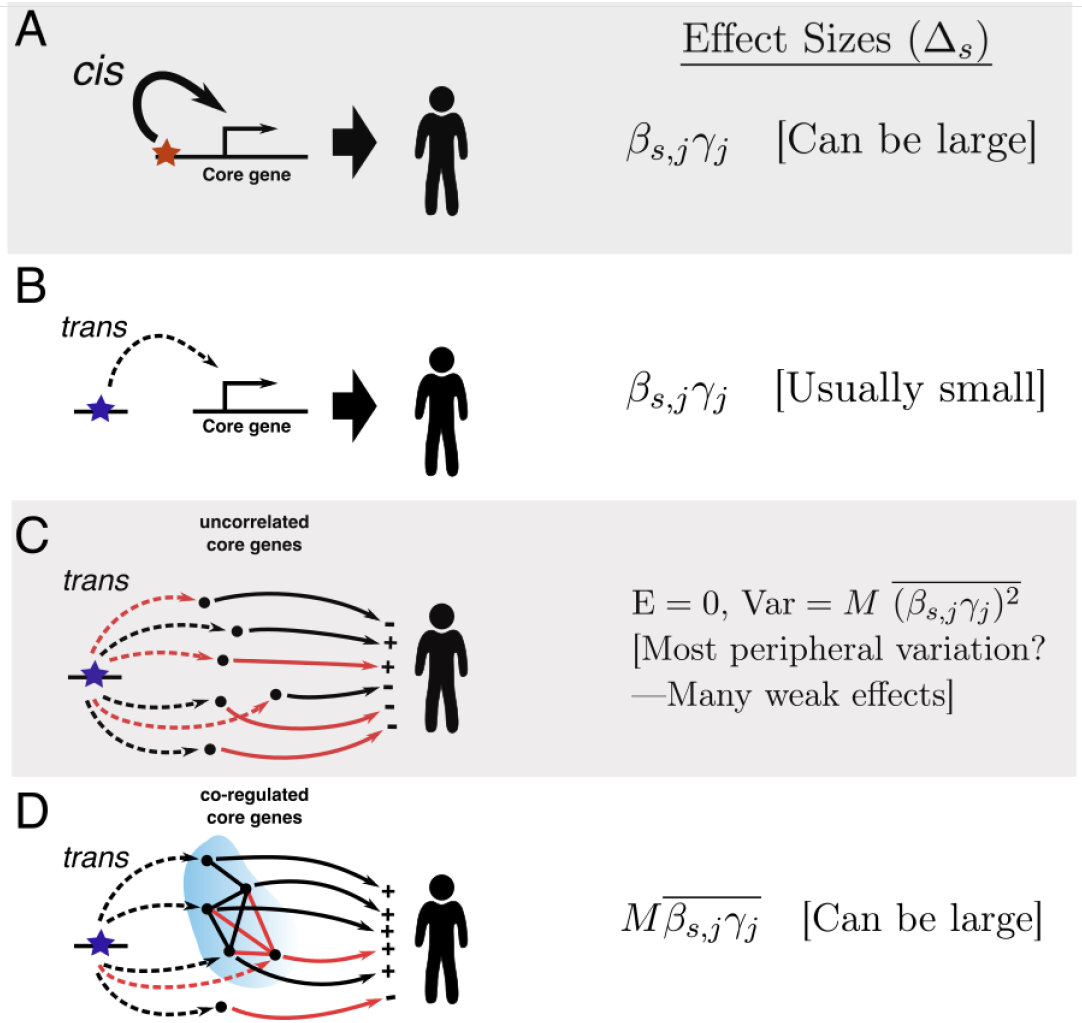
Effect sizes of cis- and trans-regulatory variants on a trait. Here the β_s_ are eQTL effect sizes of SNPs on core genes, and the γ^s^ are effect sizes of core genes on the phenotype. **A.** and **B**. For a single core gene, cis-regulatory variants will tend to have larger effect sizes on the trait compared to trans (peripheral) variants, since cis-eQTLs tend to be much stronger than trans-eQTLs. **C.** Trans-acting variants that affect many core genes will usually (but not always) have small effect sizes on the trait if the directions of effects on core genes are uncorrelated. D. Trans-regulators can have large effects on a trait if they act on many core genes in a correlated manner.(Black and red arrows indicate positive and negative effects, respectively. ‘+’ and ‘-’ indicate the sign of β_s,j_ γ_j_ for each core gene.)

### SNP effect sizes

Suppose that SNP *s* is an eQTL for core gene *j*. As before, *β_s,j_* is the effect size of SNP *s* on expression of gene *j* (each additional copy of the alternate allele at Δ*s* increases expression of *j* by *β_s,j_* units). We denote the expected change in phenotype *Y* due to one additional copy of the alternate allele as _*s*_. Suppose that gene *j* is the only core gene for which *s* is an eQTL. Then the effect size of *s*on phenotype *Y* is Δ_*s*_ = *β_s,j_ γ_j_*. Since trans-eQTLs tend to have very small effect sizes, we can expect that Δ_*s*_ will tend to be very small if *s* is in trans to *j*, compared to when *s* is in cis.

Next, what happens if *s* is a trans-QTL for multiple core genes? Now, the total phenotypic effect of *s* is determined as a sum of trans effects as mediated through each core gene *j*:

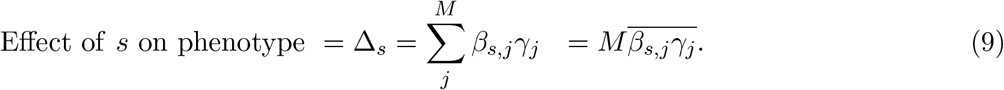

First, consider a regulatory variant that affects multiple core genes, but not in a coordinated way. In other words, the effects of SNP *s*, as mediated through different core genes may be both trait-increasing, and trait-decreasing. Specifically, if we assume that *β_s,j_ γ_j_* has an expected value of 0 and is uncorrelated across *j* then

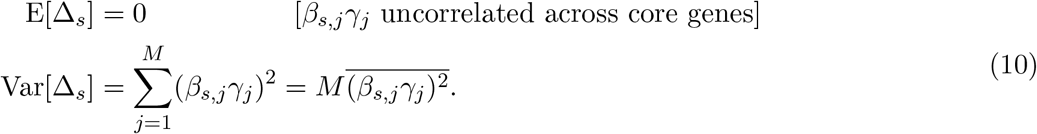

Although the effects tend to cancel out on average, the variance of the phenotypic effects scales with *M*. Although not shown here, any correlations in *β_s,j_ γ_j_* among core genes would further increase the variance.

In summary, while most SNPs would have effect sizes near zero in this model, some SNPs may have appreciable effect sizes if a preponderance of the *β_s,j_ γ_j_* happen to share the same direction of effect by chance. We hypothesize that the bulk of complex trait heritability is driven by weak random effects of this type from peripheral genes.

### Peripheral Master Regulators

In some cases, the lead hits from GWAS studies do not tag core genes, but master regulators such as KLF14 (diabetes) and IRX3/5 at the FTO locus (obesity) [31, 2]. Given that individual trans-eQTLs tend to be very weak, it seems likely that these genes drive coordinated effects on many downstream target core genes, such that the sign of *β_s,j_ γ_j_* for a given SNP tends to be systematically positive (or negative). In this case, the effect of SNP *s* is given by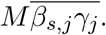 If *β_s,j_ γ_j_* tends to have the same sign across different core genes (*j*), this may potentially add up to a relatively large effect (Figure 5D).

One recent study suggests that this pattern may be a common disease architecture. Reshef *et al*., (2018) found a number of transcription factor-disease pairs for which SNPs in the transcription factor binding sites showed a persistent directional effect such that the alleles that increase binding tend to increase (or alternatively, to decrease) disease risk [48]. We interpret this as implying that increased binding of the transcription factor tends to drive directional effects on disease risk across many target genes. Thus a single variant that affects the protein or expression of the transcription factor may have a coordinated effect on many target genes.

### Diseases mediated through multiple tissues

Many traits are affected by distinct biological processes acting in different tissues. For simplicity, we largely ignore this point in the present paper. However this is relatively easy to model by adding tissue-specific subscripts to the *β*_s_ and *γ*_s_. Then, ignoring the possibility of tissue-interaction terms, the GWAS effect size on SNP *s* is the sum of tissue-specific effect sizes.

### Pleiotropy

Lastly, this model suggests a conceptual framework for interpreting variants that affect multiple traits (Box 3; Figure 6) [49, 50, 51, 52].

**Figure 6:**
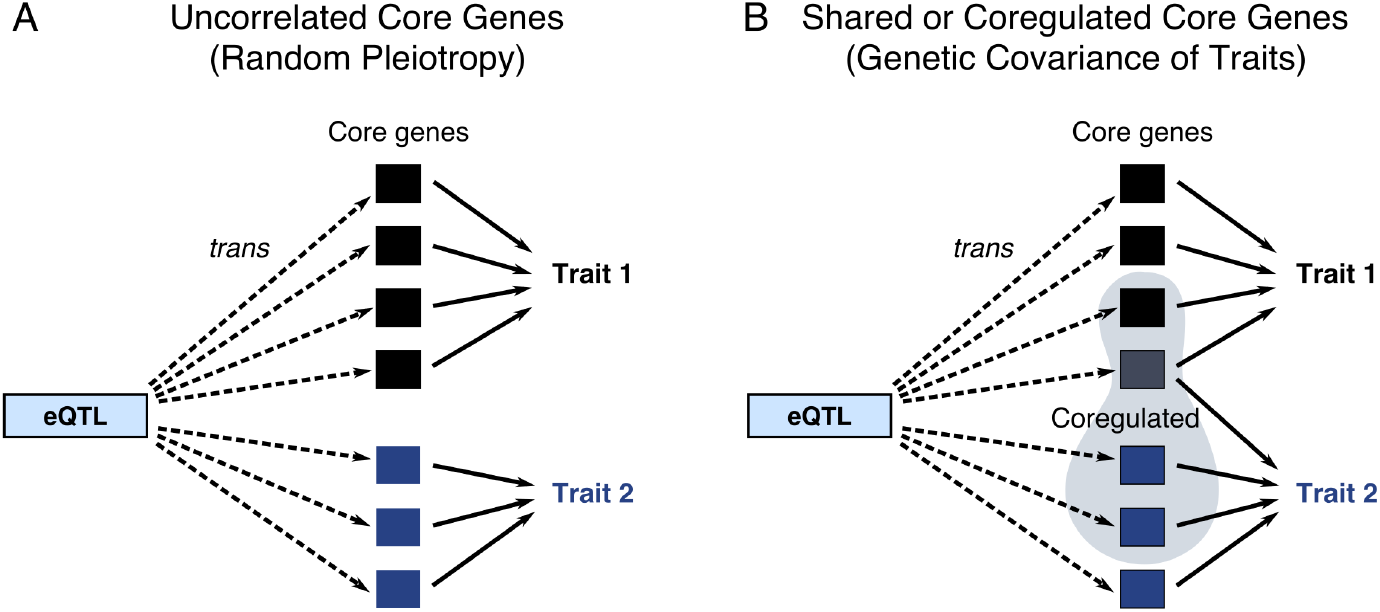
Pleiotropy and genetic correlation. **(A)** If the core genes for two traits are uncorrelated, then variants that are trans-eQTLs may affect both traits, but with uncorrelated directions of effect. **(B)** If some of the core genes are shared between traits or expression of the core genes is genetically correlated, then this may lead to genetic covariance of the traits. Genetic covariance of the traits occurs if the directions of trans-regulation and effect sizes tend to line up between the two traits in a coordinated way (i.e., that sums of γ_j,A_ γ_j,B_ for shared core genes and γ_j,A_ γ_k,B_C(j,k) across pairs of core genes, are either substantially positive, or negative overall).

First, suppose that two traits have core genes in different parts of the network (i.e., that there is no genetic covariance in the expression of the core genes). In this case, individual variants may affect both traits in a sporadic fashion: Δ_*s*_ for both traits is nonzero but with the direction of effects uncorrelated (see e.g., Figure 5C). We previously referred to these random effects as “network pleiotropy” [10] and this is related to the concept of “Type 1 Pleiotropy” [53].

Second, suppose that two traits either share core genes, or both traits have core genes in the same co-regulated networks. In these cases, the two traits will have correlated effects (i.e., genetic correlation [51, 54]) if the directions of effects tend to line up across core genes. Suppose that *γ_j,A_* and *γ_k,B_* measure the effects of expression of genes *j* and *k* on traits *A*, and *B*, respectively. (This notation simply extends the previous *γ_j_* notation to multiple traits.) Then the genetic covariance will be nonzero if the directions of the gene effects tend to line up in a consistent way, as follows (Box 3). For shared core genes we simply need the product *γ_j,A_ γ_j,B_* to tend to be consistently positive, or consistently negative. Similarly, co-regulated core genes would need to have consistently shared (or consistently opposite) directions of effects and co-regulation: i.e., that the sum of *γ_jA_ γ_kB_C*(*j,k*) across all pairs of core genes is substantially nonzero. These conditions may be met if traits are driven by overlapping genes or gene networks (as seems to be the case for psychiatric diseases [45, 55]). More trivially, this is almost guaranteed to occur if one trait contributes causally to another, downstream of genetic effects–for example, lipid levels contribute causally to coronary artery disease [52].

#### Optional Box 3.

Pleiotropy and genetic covariance of traits. Consider two traits *A* and *B*, where *Y_i,A_* and *Y_i,B_* denote the phenotypes of individual *i*, and where *M_A_* and *M_B_* denote the sets of core genes for each trait, respectively. From Eq. 1, the phenotypic covariance of these traits is

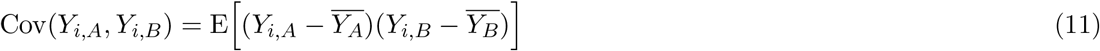

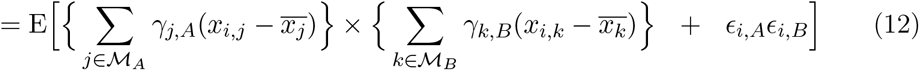

The genetic component of the covariance then depends on a sum of terms due to core genes shared between the traits, and a sum of terms based on genetic covariance of all pairs of core genes. Specifically, the genetic covariance of traits *A* and *B* is

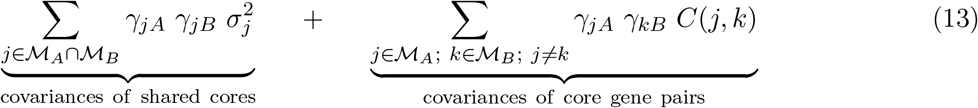

Here the first sum indexes over shared core genes, and will contribute positive trait covariance if the core genes tend to have the same directions of effects on both traits. The other sum indexes over pairs of core genes, and will contribute positive trait covariance if core gene pairs with positive expression covariance tend to have same-direction effects on both traits (and negatively correlated core genes tend to have opposite-direction effects). Reversal of these conditions would produce negative trait covariances.

## 5. Discussion

The field of human genetics has made significant strides toward elucidating the genetic basis of a wide range of complex traits. However there has been a paucity of new conceptual models for the links between genetic and phenotypic variation. In particular, how should we understand the observations that: (1) an enormous number of variants, spread widely across most of the genome affect any given trait, and that (2) together, the biggest GWAS hits generally contribute just a small fraction of the total heritability?

In this paper, our main goal was to flesh out details of the omnigenic model that we proposed last year, and to explore the implications for understanding complex trait architecture. The key points are as follows:

- Our model partitions genes into core genes (i.e., those with direct effects on the phenotype in question), and peripheral genes (non-core genes that are expressed in disease-relevant tissues). Our model suggests that peripheral genes, which can only affect the trait by modulating expression levels of core genes, are responsible for most trait heritability (Figures 1 and 2). We proposed an equation that relates expression of the core genes to the expected phenotype value (Equation 1).
- An essential component of this model is that trans-eQTLs from peripheral genes can have a big cumulative effect on the expression of core genes. The literature indicates that most of the heritability of gene expression (*~*70%) is controlled in trans (Table 1), and yet individual trans effects are almost uniformly tiny (Figure 3). This implies that expression of a typical gene is affected by huge numbers of trans-eQTLs, although we currently know little about the structure of trans-eQTL networks in humans. If we hypothesize that there are hundreds of core genes for typical disease phenotypes, each with many trans-eQTLs, these observations may start to explain why such a large part of the genome is implicated for any given trait.
- Equation 3 suggests predictions regarding the effect sizes of regulatory variants (Figure 5). Since cis-eQTLs usually have much larger effect sizes than trans-eQTLs, we may expect that many of the biggest signals in GWAS studies are cis-regulators of core genes. Second, in some cases peripheral gene-regulatory variants may become notable hits, presumably because they are trans-eQTLs for many core genes with correlated directions of effect. Third, we hypothesize that the bulk of trait heritability is driven by a huge number of peripheral variants that are weak trans-eQTLs for core genes.
- This model allows us to predict the fraction of heritability that is mediated directly through core genes, vs. through trans effects (Figure 4). If the regulation of core genes tends to be uncorrelated, then the core gene heritability simply matches the fraction of heritability that is due to cis-regulatory variants–i.e., *~*30%. In contrast, if core genes are often co-regulated, with shared directions of effects– as seems likely to us–then nearly all heritability would act through trans effects. This model may explain why even traits such as lipid levels–which are less complex than many disease phenotypes–are affected by much of the genome [8].
- Lastly, the model can also describe pleiotropic effects between different traits (Figure 6). Even for unrelated traits, it is likely that a large fraction of variants may have small effects on both traits, but with uncorrelated directions of effects. For traits that share core genes, or for which some of the core genes are in the same co-regulated networks, we can expect genetic correlation if the products γ_*j,A*_ γ_k,B_ for shared core genes and *γ_j,A_ γ_k,B_C*(*j,k*) across pairs of core genes are substantially positive, or negative, on average.

While our model is both an abstraction and a simplification of complex trait architectures, it may be helpful to interpret this model in the light of well-studied traits.

### Core genes in example traits

Some of the best-understood examples of core genes come from studies of plasma lipid levels (LDL, HDL, and triglyceride levels), which are important risk factors for heart disease. The genetics of lipid levels include both monogenic syndromes (collectively referred to as dyslipidemias) and a polygenic component, in which the latter drives most of the population-level variance. At least nine genes are currently implicated in familial hypercholesterolemia, and additional genes cause other forms of dyslipidemia [20]. The monogenic syndrome genes are closely involved in aspects of lipid metabolism or regulation and should likely be considered core genes for these traits. For example, APOB encodes Apolipoprotein B, the primary protein in LDL particles. The LDL-R protein is a receptor for LDL particles, removing them from the bloodstream and transporting them into cells–thus reducing plasma levels of LDL. Presumably additional core genes have not yet been identified as such.

Notably, most of the dyslipidemia genes are also linked to GWAS signals, indicating that common variants at these loci also contribute to lipid levels [56, 57, 58, 38, 59]. For example, 7 out of 10 genes associated with monogenic disorders of LDL-cholesterol levels are within the set of 57 genome-wide significant hit regions from a GWAS of LDL levels [20, 57].

However, while the genome-wide significant hits are highly enriched with putative core genes for this trait, it is striking that they are responsible for only a modest fraction of the heritability of LDL levels. Together, these 57 genome-wide significant loci explain *~*20% of the heritability, while all variation together explains *~*80% [5]. One study estimated that 54% of 1MB windows in the genome contribute to the heritability of extreme lipid levels [8]. Thus, in the case of LDL levels, we have clear evidence for involvement of core genes, yet they contribute only a small fraction of the genetic variance in the trait. We predict that much of the remaining variance is due to the combined contributions of many small trans effects being funneled through the core genes.

Furthermore, high LDL-cholesterol and triglyceride levels are important causal factors for coronary artery disease (CAD) and heart attack [60]. So we should expect core genes for lipids also to be core genes for CAD. Indeed, many of the key LDL and triglyceride genes do show clear involvement in CAD risk [61]. Genes involved in other physiological pathways including blood pressure, inflammation, and proliferation and repair of arterial cells are also major contributors to CAD risk [60]. This illustrates the principle that a single disease phenotype is often influenced through multiple pathways and core gene sets acting in different tissues.

However in most diseases it is currently much harder to enumerate likely core genes. In part this is because most complex diseases are poorly understood compared to lipid levels. But more fundamentally, many diseases likely have much larger core gene sets, potentially affecting multiple biological mechanisms, and potentially in multiple tissue types.

The fact that traits vary substantially in the extent of polygenicity (for example schizophrenia is substantially more polygenic than lipid levels [8, 5]) likely indicates that different traits vary greatly in the numbers of core genes and the numbers of biological processes affected. Recent work on educational attainment provides an extreme version of this phenomenon [62]. The measured phenotype of educational attainment is affected by many different aspects of psychology and health; presumably each of these has its own core genes, and the measured effect sizes on each SNP are weighted averages across all these simpler traits.

Lastly, it is important to note that our definition of core genes is a simplification of a more-complex reality. There are various edge cases that are hard to classify. For example, PCSK9, which is an important drug target for lipid levels, acts by degrading LDL Receptor proteins. It is tempting to label this as a core gene, though strictly speaking it acts through protein regulation of the LDLR gene and by our definition should thus be considered peripheral. As another example, many receptor genes are involved in *receiving* extra-cellular signals such as hormones or cytokines, and then driving internal cellular regulatory networks.

We are inclined to regard these as potential core genes as they interact directly with external signals, leading to changes in cellular function; however they do not fit neatly within our definition.

### Peripheral master regulators

Since trans-eQTL effect sizes tend to be extremely small, most peripheral genes have individually small effects on traits. But there are now several examples of variants that likely affect many core genes in a coordinated way, and thus stand out as important GWAS hits (Figure 1C).

For example, a variant at the KLF14 locus is associated with dyslipidemia, insulin dependence, and type 2 diabetes [31]. This variant, which is a cis-eQTL for KLF14, is a trans-eQTL for a network of 385 other genes in adipose tissue. Several of the target genes are strong candidates for driving aspects of the organism-level phenotypes, and it is likely that the overall effects of KLF14 is mediated through multiple loci (i.e., core genes) in this network.

Similarly, a SNP in the FTO gene that is a cis-eQTL for IRX3 and IRX5, is associated with triglyceride levels, obesity and diabetes [57, 63, 2]. These two alleles control the fractions of adipocyte precursors that differentiate into white and beige adipocytes respectively [2]. In both the KLF14 and FTO examples, the SNPs alter transcriptional programs with downstream consequences on disease risk.

As a third example, circadian rhythms are controlled by a well-understood set of transcriptional regulators and repressors that drive daily cycling of thousands of genes [64, 65]. A recent GWAS for whether people are “morning people” or “evening people” identified 351 loci, with strong enrichment of signal among genes expressed in the brain and pituitary [66]. Notably, the peaks included nearly all of the key circadian regulators. In our terminology these are not core genes as they do not exert direct causal effects on chronotype, but instead act as coordinated master regulators of many downstream core genes that drive daily physiological cycling.

We anticipate that many of the examples of transcription factors, chromatin modifiers, and other regulatory genes that have emerged as strong hits in disease studies act as peripheral master regulators, driving coordinated regulation of many core genes. Such genes are of particular interest for understanding biological drivers of a trait, and in some cases stand out as lead GWAS hits, although as we discuss next, GWAS studies may actually be biased against finding such variants.

### The role of selective constraint in shaping the heritability landscape

Our main goal in this paper has been to understand how existing human genetic variation combines with gene regulatory architecture to produce the observed distribution of heritability across the genome. But it is important to note that the landscape itself is an evolved property that has been strongly shaped by the action of purifying selection. The strength of purifying selection against a causal variant increases rapidly with its effect sizes, both on the trait in question and through pleiotropic effects of that variant on other traits [67]. This means that the allele frequencies for trait-associated variants tend to be inversely related to their effect sizes, and that larger-effect size variants generally contribute little to heritability [68].

There are two important implications. First, the contributions of different genes to heritability is unlikely to be proportional to their intrinsic biological importance for the trait in question. Specifically we can expect that selective constraints tend to flatten out the landscape of heritability such that the most important genes contribute less than might be expected given their intrinsic importance.

Second, the strength of pleiotropic effects may be particularly strong for some genes, such as master regulators, and thus the contributions of these genes to heritability may be greatly reduced compared to their intrinsic importance. These points were discussed recently in a paper showing differences between yeast mapping results when using natural polymorphisms versus gene knockouts [32].

### Next steps in deciphering complex traits

Broadly speaking, genetic studies of complex traits can make two kinds of contributions: (1) prediction of individuals at risk of disease, and (2) elucidation of biological mechanisms and potentially therapeutic targets.

With recent progress on polygenic risk scores, the GWAS field is now making meaningful strides toward the goal of risk prediction in clinical applications [9, 69]. Accurate polygenic risk prediction depends on having accurate estimates of tiny effect sizes across millions of SNPs. Polygenic prediction can be done without a deep understanding of biological mechanisms of disease, but it does require enormous sample sizes. Therefore, to achieve the full potential of polygenic prediction, it will be essential to continue building larger GWAS samples for the major diseases. Fortunately, the cost and difficulty of building large GWAS samples continue to drop, as a result of cheaper genotyping, the emergence of large public biobanks in multiple countries, and the growing use of genotyping in health care and in personalized genomics companies. We fully support these efforts.

A more difficult question is to determine the best paths forward for linking GWAS data to biological mechanism. In our view, the current biggest gap is the very limited knowledge of trans-regulatory networks. If we had high quality trans-regulatory networks and trans-QTL information, then this could potentially be combined with GWAS effect size estimates to enable a complete description of core and peripheral genes, and the flow of genetic effects through the regulatory network. Existing methods that combine GWAS and eQTL data, such as PrediXcan and TWAS, provide a roadmap as to how we might conceptualize this analysis, but these methods are currently limited by our very poor knowledge of trans-QTLs [70, 71]. But with high-quality network information, it may be possible to extend this concept to perform joint inference on all genes to identify which genes are core genes, which genes are master regulators and which are weaker peripheral genes.

The key question then is how to infer regulatory networks. One approach is through trans-eQTL mapping, but this requires extremely large sample sizes. Studies of whole blood are starting to approach the required sample sizes, but extremely large samples are far less practical for most other tissues or cell types. Alternatively we are optimistic that high-throughput experimental perturbation methods may help to fill this gap [72, 73, 74]. In brief, these approaches work by perturbing one or more genes, and then measuring the effects on expression of other genes. A natural hypothesis is that a cis-eQTL variant for a particular gene would recapitulate the same directions of effect as the experimental perturbation. These types of approaches are still in their infancy but are promising as they are far more scalable than trans-eQTL mapping.

Another open question is the value of deep sequencing to identify rare variants of larger effects. These approaches have so far had mixed success, depending on the disease [75, 76, 77, 78]. In principle, rare variants of larger effect can provide orthogonal information to the common variant signal, should generally be more proximate to the mechanism of action, and may help to identify important genes that are refractory to common variation. On the other hand, most of these studies continue to be underpowered at current sample sizes. As sequencing costs continue to drop, we believe that deep sequencing will continue to be an important tool that provides complementary information, while recognizing that it is no panacea. Ultimately a full mechanistic dissection of complex traits will require a combination of all of these kinds of approaches, along with detailed functional biology of key targets.

## Summary

This paper aims to provide a simple, but formal, model for the links between genetic variation, expression of core genes, and disease risk. We have argued previously that most of the heritability for typical complex traits is mediated through genes that have only distant connections to disease biology. Here we have expanded on this theme, proposing that this is a consequence of known features of cis- and trans-eQTL architecture.

## Acknowledgements

We thank many people for helpful conversations or comments including Evan Boyle, Diego Calderon, Jake Freimer, Ziyue Gao, Arbel Harpak, Mark McCarthy, Hanna Ollila, Luke O’Connor, Molly Przeworski, Andrey Rzhetsky, Guy Sella, Eilon Sharon, Gavin Sherlock, Yuval Simons, and Nasa Sinnott-Armstrong. This work was supported by NIH grants HG008140, and HG009431.

## Footnotes

1 As defined in this paper, *omnigenic* has a more precise meaning than the term *polygenic*. *Polygenic* can be used to describe the involvement of anything from tens of loci to every variant in the genome and would include *omnigenic* as a special case, toward the high end of the polygenic spectrum. We also use the term *omnigenic model* to refer to our model of complex trait architecture in which heritability is mainly driven by peripheral genes that trans-regulate core genes.

2 It is also worth distinguishing our model from Fisher’s classic infinitesimal model [29, 30]. The infinitesimal model was originally developed in the pre-molecular era. While fundamentally important for understanding patterns of inheritance, it does not tell us how many causal variants to expect in practice, nor about the molecular mechanisms linking variation to phenotypes.

3 This definition of core genes is narrower and more precise than used in our original paper [10].

4 Here, regulatory networks would include diverse aspects of regulation of core genes by other gene products within cells, including regulation of mRNA or protein expression levels and transcript usage, post-translational modifications, and protein localization. We exclude extra-cellular signaling such as hormones or cytokines from this definition, so that signaling receptors can be core genes (see the Discussion). Notice that core genes and peripheral master regulators are defined with respect to a given trait or disease. Furthermore, these definitions depend on gene functions and regulatory architecture, and do not depend on the presence of mappable variants.

5 The multi-tissue extension of this model can be handled by adding tissue subscripts to the γ_s_ and *x*s but does not change the overall conclusions and is not considered further.

6 This model is described in terms of quantitative phenotypes. Presence or absence of a disease is often modeled by assuming that disease risk is determined by an underlying quantitative liability scale.

7 We assume that trans heritability of core genes is similar to average trans heritability. If core genes are under particularly strong purifying selection, then we may expect the cis variance to be reduced and hence the fraction of trans heritability for core genes would, if anything, be higher.

8 The third possibility, that the average of *γ_j_ γ_k_C*(*j, k*) is substantially negative, is mathematically possible but seems less biologically relevant as it requires a preponderance of gene pairs with configurations such as anti-correlated expression but shared directional effects.

9 This calculation assumes that γ^2^_*j*_ is independent of *σ*^2^_*j*,cis_ and *σ*^2^_*j*_,trans. If instead, γ^2^_*j*_ is negatively correlated with *σ*^2^_*j*,cis_, then this would further reduce the cis heritability of *Y*.

10 Note that some variants with trans effects on core gene *j* may be in cis to another core gene, *k*, say. In that case, the transeffect on gene *j* acts through the regulatory network, effectively like a peripheral variant, although the GWAS signal itself would be found in cis to *k*. This effect would slightly increase the measured contribution of core gene variants to heritability, although it seems likely to be a modest effect unless core genes make up a large fraction of all genes.

